# Cathepsin K as a Key Protease in Processing of SARS-CoV-2 Spike Activation Sites and a Target of Dual-Inhibition

**DOI:** 10.1101/2025.10.03.680348

**Authors:** S. Yasin Tabatabaei Dakhili, Dipon Saha, Preety Panwar, Eliot Mar, Olivier Hinse, Yu Seby Chen, Bryan J. Fraser, Cheryl Arrowsmith, Masahiro Niikura, Filip Van Petegem, Dieter Brömme

**Author notes:** authors contributed equally (S. Yasin Tabatabaei Dakhili, Dipon Saha).

## Abstract

SARS-CoV-2 relies on host proteases to prime its spike protein for cell entry through either the endosomal or plasma membrane pathway. Although cysteine cathepsins are known to mediate the endosomal route, the identity of the dominant enzyme has remained unclear. Here, we identify human Cathepsin K (hCatK), a lysosomal cysteine protease, as a previously unrecognized yet functionally important mediator of spike activation. While human Cathepsin L (hCatL) has long been regarded as the principal endosomal protease for spike processing, inhibition of hCatK with the selective inhibitor Odanacatib suppressed viral infection in endothelial cells as effectively as the broad-spectrum cysteine protease inhibitor E-64d, implicating hCatK as a key driver of spike processing during the endosomal viral entry. Comprehensive enzymatic profiling demonstrated that hCatK exhibits 24- to 63-fold higher catalytic efficiency toward the Furin-cleavage site (FCS) sequence than hCatL and displays a distinct substrate-recognition pattern at the Omicron FCS relative to the Wuhan variant. We further demonstrate that hCatK is an off-target of Nirmatrelvir, a clinically approved 3CL-Mpro inhibitor, with a sub-micromolar potency (IC_50_ = 0.6 ± 0.1 µM). A 1.9 Å crystal structure of the hCatK–Nirmatrelvir complex delineates the molecular basis of inhibitor binding and supports the rational design of dual-acting antivirals. Collectively, these findings redefine the landscape of host proteases involved in SARS-CoV-2 spike activation and establish hCatK as a previously overlooked but strategic target for antiviral intervention.

## Introduction

Host proteases are critical for SARS-CoV-2 entry^1^. They process the spike protein at defined sites-TMPRSS2 cleavage site (TCS) and FCS to enable membrane fusion. Most studies have focused on cell-surface proteases such as Furin and TMPRSS2, which mediate spike activation at the plasma membrane^2,3^. However, SARS-CoV-2 can also enter cells via an endosomal route. This cathepsin-dependent endosomal entry pathway is increasingly utilized by emerging viral variants but remains incompletely understood at the molecular level^4,5^. Although hCatL and B have been implicated in spike activation during endosomal entry, much of the supporting evidence relies on broad-spectrum cysteine protease inhibitors and lacks kinetic characterization, or mapping of cleavage site^6,7^. As a result, the individual contributions of other cathepsins, such as hCatK and hCatS, lysosomal cysteine proteases expressed in lung, vascular, and endothelial cells, remain poorly define^8-10^. In this regard, hCatK, has emerged as a notable candidate, as its selective inhibitor ONO-5334 was identified among the top compounds with antiviral activity against SARS-CoV-2 in a large-scale drug-repurposing screen^11^. Addressing these gaps are essential to establish mechanistic evidence of spike activation by specific cathepsins and for the design of therapeutics that target both viral and host enzymatic components^12,13^.

Here, we used Odanacatib, a selective hCatK inhibitor, to confirm the direct role of this enzyme in SARS-CoV-2 infection of endothelial cells^14,15^. To further elucidate the mechanistic role of cysteine cathepsins in spike processing, we compared the kinetic efficacy of selected cysteine cathepsins and canonical convertases (Furin and TMPRSS2) on TCS and FCS-related spike peptides and determined the cleavage site pattern by LC-MS/MS analysis.

Given the sequence similarity between the 3CL-Mpro substrate motif (`AVLQ-SGFR`) and known cathepsin recognition sequences, we hypothesized that cysteine cathepsins, particularly hCatK, might share overlapping substrate preferences with the viral protease^16-18^. To test this hypothesis, we performed kinetic analyses of selected cysteine cathepsins using the 3CL-Mpro substrate and subsequently determined the crystal structure of hCatK in complex with Nirmatrelvir, an FDA-approved 3CL-Mpro inhibitor^19^. This approach allowed us to assess potential cross-reactivity and define the structural basis for dual inhibition of both viral and host proteases.

Together, these integrated biochemical, kinetic, and structural approaches aim to redefine the molecular framework of spike activation within the endosomal pathway and uncover hCatK as a previously unrecognized but therapeutically actionable host protease in SARS-CoV-2 infection.

## Materials and Methods

FRET-based peptides, Ac-`AE` (Edans) QTNSPRRARSVK (Dabcyl) (FCS-Wu), Ac-`AE` (Edans) QTKSHRRARSVK (Dabcyl) (FCS-Omic), and Ac-`AE` (Edans) LPDPSKPSKRSFIK (Dabcyl) (TSC), were custom synthesized (>98% purity) by Biomatik Co., Ontario, Canada. Recombinant SARS-CoV-2 Spike proteins S1 and S2 were purchased from RayBiotech Life Inc. (GA, USA). Human CatK antibody (Abcam; AB37259) and Alexa Fluor dye 488 (Abcam; AB112125) were obtained from Abcam (Cambridge, UK), and viral nucleocapsid protein antibody (GTX635674) from GeneTax (Irvine, CA). Amicon^®^ Ultra - 15 Centrifugal Filters were from Merck Millipore Ltd, Irland). Human CatK was expressed in *Pichia pastoris* and purified as described in and human CatS was expressed in *E. coli* BL21(DE3)^20,21^. Recombinant TMPRSS2 (dasTMPRSS2) was produced and purified as previously described^22^. Human recombinant CatB had been kindly provided by J. Mort^†^ from the Shriner’s Hospital for Sick Children (Montreal, QC, Canada). Recombinant human Furin and human liver CatL were obtained from Fisher Scientific, Canada. Furin Inhibitor I (Decanoyl-`RVKR`-CMK) was obtained from EDM Millipore Corp., USA, L-3-carboxy-*trans*-2–3-epoxypropionyl-leucylamido-(4-guanidino)-butane ethyl ester (E-64d) from Sigma-Aldrich (Sigma-Aldrich Canada, Oakville, ON, Canada), Odanacatib, Camostat from Selleck Chemicals (Houston, TX), and Nirmatrelvir from A2B Chem, LLC (San Diego, CA).

### SARS-CoV-2 virus propagation

SARS-related Coronavirus 2 (SARS-CoV-2), isolate USA-WA1/2020, was obtained from BEI Resources (Cat# NR-52281) and used in all experiments. All work with SARS-CoV-2 was conducted in BSL-3, a Containment Level 3 facility at Simon Fraser University, following approved protocols (SFU Biosafety Committee Permit #361-2022 and #361-2023). The stock virus was produced by amplifying the initial seed virus in VeroE6/TMPRSS2 cells (JCRB Cell Bank, Cat# JCRB1819) through two passages before use in experiments.

### Endothelial cell culture and virus infection

EA.hy 926 endothelial cells (ATCC, Burlington, ON) were seeded at a density of 50,000 cells per well in Dulbecco’s Modified Eagle Medium (DMEM) containing 10% fetal bovine serum in a multi-well plate and infected with 50,000 pfu of the virus in 50 µL of culture medium added to the existing medium and incubated at 37°C for 48 hours at 5% CO_2_ in the presence or absence of E-64d, Camostat, Odanacatib or Nirmatrelvir (0 to 20 µM). Following treatment and removing the medium, cells were scraped into 200 µL of RIPA containing a proteinase inhibitor cocktail (ThermoFisher Scientific, Canada) and incubated for 10 min on ice with occasional vortex mixing for the Western blot samples. The samples were boiled before being removed from BSL-3. For microscopic studies, cells were fixed in 0.2% glutaraldehyde and 4% formalin (1 mL per well) for 30 min at room temperature after removing the medium. The plates were disinfected and removed from BSL-3 for staining.

Immunostaining for hCatK was performed using a protocol optimized for fixed cell imaging. In parallel, viral nucleocapsid protein staining was carried out using a specific antibody to detect infection levels. Briefly, 0.2% glutaraldehyde and 4% formalin fixed EA.hy 926 cells were permeabilized using 0.1% Triton X-100. After blocking with 1% bovine serum albumin (BSA), cells were incubated with primary antibodies against CatK and viral nucleocapsid protein, followed by fluorescently labeled secondary antibodies (Alexa Fluor 647; ThermoFisher Scientific, Canada). Actin cytoskeleton was stained using Phalloidin conjugated to dye (Alexa Fluor 488), and nuclei were counterstained with DAPI. Imaging was performed using a fluorescence microscope (Olympus BX61 Fluorescence microscope, Olympus, USA).

### 3CL-Mpro protein expression and purification

Hisx6-tagged WT-3CL-Mpro (expression vector kindly provided by Dr. C.M. Overall, University of British Columbia, UBC) was expressed in *E. coli* BL21-DE3. The inoculated culture (2–6 L) was grown in LB broth medium at 37°C in the presence of 100 microgram/L ampicillin till OD_600nm_ reached 0.6. The temperature was then lowered to 16°C, and expression was induced overnight with 0.5 mM IPTG. The cells were harvested by centrifugation at 5,000g at 15°C for 10 min in an Avanti J-E centrifuge (Beckman Coulter, Inc) and resuspended in a lysis buffer (50 mM Tris [pH 7.5], 300 mM NaCl). The cells were lysed by sonication on ice and centrifuged at 20,000g for 30 min at 4°C. The supernatant was loaded on a Ni-NTA column previously equilibrated with a lysis buffer at RT. The His-tagged 3CL-Mpro enzyme was eluted from the column in 50 mM Tris, pH 7.5, 300 mM NaCl, 300 mM imidazole. The elution was buffer exchanged to 50 mM Tris, pH 7.5, 150 mM NaCl overnight in presence of PreScession protease to get rid of imidazole and His-tag. Then the enzyme was purified using the Ni-NTA column and finally, loaded onto a Superdex 200 size-exclusion column (GE Healthcare) using an AKTA purifier core system (GE Healthcare). The column was pre-equilibrated with a degassed filtered buffer (50 mM Tris [pH 7.5], 150 mM NaCl). The final protein was collected, concentrated and kept in −80 °C for future use. To express Hisx6-tagged-PreScission protease (expression plasmid kindly provided by Dr. Petegem, UBC), the plasmid was inoculated with *E. coli* BL21-DE3 culture (2–6 L) and grown in LB broth medium at 37°C in the presence of 100 µg/L ampicillin till OD_600nm_ reach 0.6. The temperature was then lowered to 16°C, and expression was induced overnight with 0.5 mM IPTG. The cells were harvested by centrifugation at 5,000g at 15°C for 10 min in an Avanti J-E centrifuge (Beckman Coulter, Inc) and resuspended in a lysis buffer (50 mM Tris [pH 7.5], 300 mM NaCl). The cells were lysed by sonication on ice and centrifuged at 20,000g for 30 min at 4°C. The supernatant was loaded on a Ni-NTA column previously equilibrated with a lysis buffer at RT. The His-tagged PreScission enzyme was eluted from the column in 50 mM Tris, pH 7.5, 300 mM NaCl, 300 mM imidazole. Purified protein was kept at −80 °C in presence of 5mM beta-mercaptoethanol until further use.

### Enzyme kinetic assays

All enzyme assays were performed at room temperature. Cleavage of fluorogenic substrates was monitored using a BioTek Synergy H4 microplate reader. Active site titrations of cathepsins, TMPRSS2, and Furin were conducted using E-64d, Nafamostat, and Decanoyl-RVKR-CMK, respectively. To determine *k*_cat_ and K_M_ values, assays were performed at constant enzyme concentrations with increasing concentrations of FCS-Wu, FCS-Omic, and TCS substrates, respectively. Cathepsin assays were conducted in 0.1 M sodium acetate buffer (pH 5.5) containing 2.5 mM dithiothreitol (DTT) and 2.5 mM ethylenediamine-tetraacetic acid (EDTA). Cathepsins were used at final concentrations ranging from 10 to 400 nM, and substrate concentrations ranged from 0 to 90 μM. Furin assays were performed in 0.1 M HEPES buffer (pH 7.5) supplemented with 0.5% Triton X-100 and 1 mM CaCl_2_. Furin was used at a final concentration of 10 nM, and substrate concentrations ranged from 0 to 60 μM. 3CL-Mpro assays were carried out in 0.05 M Tris-HCl buffer (pH 7.5) containing 10 mM DTT. The enzyme was used at a final concentration of 100 nM, with substrate concentrations ranging from 0 to 60 μM. TMPRSS2 assays were performed in 25 mM Tris buffer (pH 8.0) containing 75 mM NaCl and 2 mM CaCl_2_ with a TMPRSS2 concentration of 12 nM, and substrate concentrations between 0 to 60 μM. All assays were performed in two independent experiments, each in triplicate. Data were plotted using the Michaelis–Menten model and analyzed via nonlinear regression in GraphPad Prism (Version 5.0, GraphPad Software, La Jolla, California, USA). Inhibition assays were conducted in the presence of increasing concentrations of Nirmatrelvir and Z-`FR`-AMC (5–30 μM) for cathepsins and MCA-`AVLQSGFRK` (Dnp) `RR` for 3CL-Mpro.

### LC–MS/MS enzyme assays and LC–MS/MS data acquisition on TIMS-TOF Pro2

FRET-based peptide substrates (50 µM) as listed above were added to the appropriate activity buffer corresponding to each enzyme. The enzyme was then added to the mixture, which was incubated at room temperature for 1 hour (CatK, S, L, B 5-400 nM; Furin 4 nM; and TMPRSS2 12 nM). The reaction was stopped by adding the corresponding inhibitor and 60% (v/v) methanol. The mixture was stored at –20 °C for 2 hours, followed by centrifugation at 21,000 × g for 10 min to remove the enzyme. Each reaction was performed in duplicate. 20 ng of peptide was injected and separated on-line using a NanoElute 2 UHPLC system (Bruker Daltonics) with Aurora Series Gen3 (CSI) analytical column, (25 cm x 75 μm 1.7 μm C18 120Å, with CSI fitting; Ion Opticks, Parkville, Victoria, Australia). The analytical column was heated to 50°C using a column toaster M (Bruker Daltonics). The NanoElute thermostat temperature was set at 7°C. Buffer A consisted of 0.1% aqueous formic acid and 0.5% acetonitrile in water, and buffer B consisted of 0.1% aqueous formic acid and 0.5% acetonitrile in water. Before each run, the analytical column was conditioned with 4 column volumes of buffer A. The analysis was performed at 0.30 μL/min flow rate. A standard 30 min gradient was run from 2% B to 12% B over 15 min, then to 33% B from 15 to 30 min, then to 95% B over 0.5 min, held at 95% B for 7.72 min.

Liquid chromatography was coupled to a Trapped Ion Mobility - Time of Flight mass spectrometer with dual TIMS (TimsTOF Pro2; Bruker Daltonics, City, State). The Captive Spray ionization source was operated at 1600 V capillary voltage, 3L/min drying gas, and 200°C drying temperature. During analysis, the TIMS-TOF Pro 2 was operated with Parallel Accumulation-Serial Fragmentation (PASEF) scan mode for DDA acquisition. The MS and MS/MS spectra were collected in positive mode, from m/z 100 Th to m/z 1700 Th, and from ion mobility range (1/ K_0_) 0.7 V*s/cm^2^ to 1.3 V*s/cm^2^. A polygon filter was applied to the mass-to-charge and the ion mobility plane to include the most likely peptide precursors and to reduce singly charged background ions.

TIMS-MS scan was set at equal ramp time and accumulation time of 100 ms, at a rate of 9.42 Hz (100% duty cycle). Active exclusion was enabled with a 0.05 min release and reconsidered if their intensity increased more than 4 times. For each TIMS cycle, 3 PASEF MS/MS scans were recorded (total cycle time 0.42 s). Target Intensity for parent ions was set to 10,000 cts/s with a threshold of 1,000 cts/s. Isolation widths were set at 2.07 m/z starting m/z 400 Th and 3.46 m/z ending m/z 1000 Th. The collision energy was ramped linearly as a function of mobility value from 27eV at 1/k0 = 0.7 V·s/cm^2^ to 55 eV at 1/k0 = 1.35 V·s/cm^2^.

### Crystallization and structural analysis of Nirmatrelvir-hCatK complexes

Recombinant hCatK was expressed in *P. pastoris* and purified as previously described.^20^ To obtain an Nirmatrelvir-hCatK complex, Nirmatrelvir was dissolved in DMSO to 50 mM and added to hCatK (1 mg/mL) in 100 mM sodium acetate pH 5.5, 2.5 mM DTT and 2.5 mM EDTA. The protein-inhibitor complex was then concentrated to 5 mg/mL using centrifugal filters with a 10 kDa molecular weight cutoff. Crystals of Nirmatrelvir-hCaK complexes were obtained by equilibrating 0.7 µL of the protein-inhibitor mixture and 0.7 µL of reservoir solution (0.96 M sodium citrate pH 7) in sitting-drop vapor diffusion system at 22 °C. Suitable crystals were formed within 3 weeks at room temperature. Crystals were fished out and incubated in a cryoprotectant solution (the cryoprotectant containing 1M sodium citrate, 25% Ethylene Glycol and 2 mM Nirmatrelvir) for 2 min before flash-frozen in liquid nitrogen for data collection.

Diffraction data from a single crystal were collected at 100 K with a wavelength of 0.98 Å using a DECTRIS EIGER2 XE 16M detector at beamline BL12-2 of the Stanford Synchrotron Radiation Lightsource (Menlo Park, California, USA). The dataset was processed with the program iMosflm version 7.2.1 and the intensities scaled with SCALA to a resolution of 1.62 Å for Nirmatrelovir-bound hCatK. Phasing was performed by molecular replacement using human wild-type CatK structure (PDB: 5TUN)^23,24^. The restraints for the complexed Nirmatrelvir were generated using ELBOW and fitted into the model during refinement^25^. All modeled structures were refined by cycles of automated refinement in PHENIX-COOT in the PHENIX program suite^26,27^. Structural images were prepared using PyMOL software (PyMOL Molecular Graphics System, Version 3 Schrödinger, LLC).

## Results

### Effect of cysteine protease inhibitors on the infection of endothelial cells with SARS-CoV-2

To understand the importance of cysteine cathepsins in endosomal entry, we used the EA.hy926 endothelial cell (EC) line as a model. This choice was based on the relatively low TMPRSS2 expression in ECs, suggesting that viral infection in these cells likely relies considerably on the endosomal pathway^28,29^. Interestingly, a comparison between SARS-CoV-2 Wuhan variant-infected ECs (Fig. 1A) and uninfected controls (Fig. 1B) revealed a 7-fold upregulation of hCatK (Fig. 1C). To investigate the potential specific role of hCatK in this process, we treated ECs with Odanacatib, a selective hCatK inhibitor, along with E-64d, a broad-spectrum cysteine cathepsin inhibitor (Fig. 1D). In addition, cells were treated with the TMPRSS2 inhibitor, Camostat^30^ and the 3CL-Mpro inhibitor, Nirmatrelvir^19^. Viral infection was measured by quantifying the percentage of ECs positive for intracellular nucleocapsid protein. All four inhibitors, E-64d, Odanacatib, Camostat, and Nirmatrelvir, reduced infection in a dose-dependent manner (Fig. 1C). Interestingly, E-64d and Odanacatib showed similar inhibitory effects, with EC_50_ values of 7.5 ± 2.2 μM and 9.0 ± 2.6 μM, respectively (Fig. 1F), suggesting that hCatK is the dominant contributor to viral cell entry among cathepsins. Camostat, a TMPRSS2 inhibitor, also showed a similar potency (EC_50_ of 6.1 ± 2.2 μM) as E-64d and Odanacatib, indicating that both the endosomal and cell-membrane entrance pathway were used to a comparable extent by SARS-CoV2 in this EC line. Nirmatrelvir, developed as a highly selective inhibitor of 3CL-Mpro, demonstrated a 3-to 4-fold greater potency in this assay with an EC_50_ of 2.2 ± 0.4 μM in preventing the proliferation of the virus. These findings suggest that hCatK plays a dominant role in mediating SARS-CoV-2 entry into ECs.

**Figure 1:**
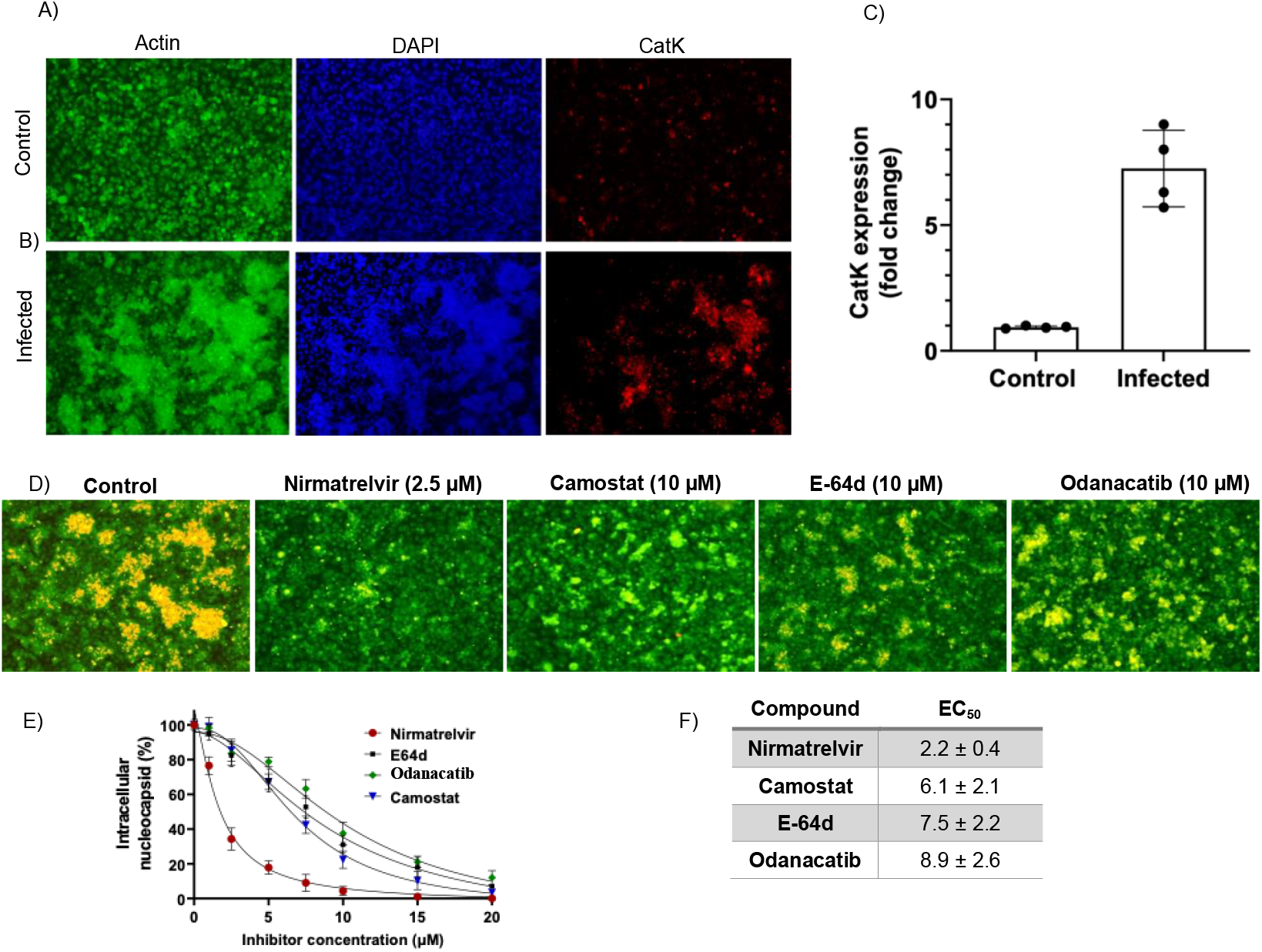
Infection-induced upregulation of CatK and antiviral effects of protease inhibitors in EA.hy926 endothelial cells. (A–B) Immunofluorescence images showing Actin (green, Phalloidin), nuclei (blue, DAPI), and CatK (red) in uninfected control (A) and infected (B) EA.hy926 cells. Infection leads to increased CatK expression and altered cytoskeletal morphology. Scale bar: 100 µm. (C) Quantification of CatK expression (fold change) in control vs. infected cells. Data are shown as mean ± SEM (n = 4); ***p < 0.001. (D) Immunostaining for viral nucleocapsid protein (yellow) and actin (green) after treatment with Nirmatrelvir (2.5 µM), Camostat (10 µM), E-64d (10 µM), and Odanacatib (10 µM). (E) Dose-dependent inhibition of intracellular nucleocapsid levels by the four compounds. Data represent mean ± SEM (n = 3). (F) EC_50_ values of the tested compounds calculated from the dose-response curves in (E).

### Kinetic analysis of Furin, TMPRSS2, and cysteine cathepsins using substrates representing processing sites in SARS-CoV-2 spike proteins

To clarify the specific contributions of cysteine cathepsins to SARS-CoV-2 Spike protein processing, we designed three Förster resonance energy transfer (FRET)-based substrates corresponding to the wild-type Wuhan FCS Furin cleavage site (FCS-Wu), the Omicron FCS (FCS-Omic), and the TMPRSS2 cleavage site (TCS) (Fig. 2A). Kinetic parameters were measured for four major human cysteine cathepsins (CatK, S, L, and B), alongside TMPRSS2 and Furin, which represent proteases related to the endosomal and plasma membrane viral entrance pathways, respectively.

**Figure 2:**
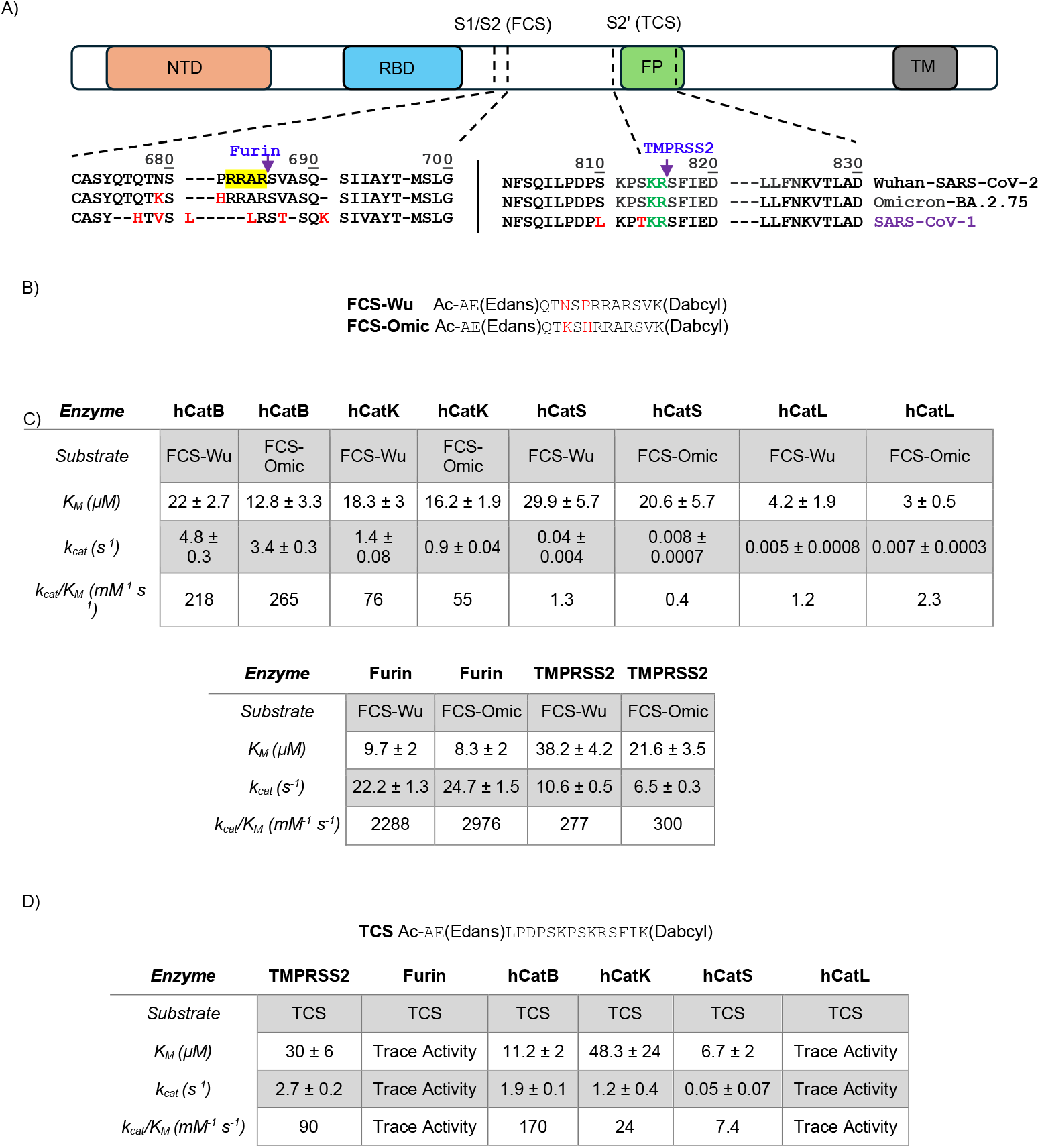
Kinetic analysis of the cleavage activities of human cathepsins and TMPRSS2 and Furin at S1/S1 and S2’ cleavage site areas in SARS-CoV-2 Spike protein. (A) Schematic representation of the Spike (S) protein, highlighting the Furin and TMPRSS2 cleavage sites. The amino acid sequences of S proteins from various SARS-related coronaviruses are aligned for comparison. (B) Table compares the kinetic parameters of selected cysteine cathepsins toward FCS-Wu and FCS-Omic substrates. (C) Table compare the kinetic parameters of Furin and TMPRSS2 using two fluorogenic substrates derived from the S1/S2 site: FCS-Wu (Ac-`AE`(Edans)QTNSPRRARSVK(Dabcyl)) and FCS-Omic (Ac-`AE`(Edans)QTKSHRRARSVK(Dabcyl)). (D) Table compare the kinetic parameters of Furin, TMPRSS2, and selected cysteine cathepsins using the synthetic fluorogenic substrate TCS (Ac-`AE`(Edans)LPDPSKPSKRSFIK(Dabcyl). The kinetic data were obtained from two independent experiments, each performed in triplicate. Values are reported as mean ± standard deviation (SD).

The results showed the highest catalytic efficiency for Furin for both FCS-Wu and FCS-Omic (k_cat_/K_M_= 2,288 and 2,976 *mM*^*-1*^ *s*^*-1*^, respectively). Human Cathepsin B (hCatB) cleaved both FCS-Wu and FCS-Omic substrates (k_cat_/K_M_ = 218 and 265 *mM*^*- 1*^ *s*^*-1*^, respectively) at levels comparable to TMPRSS2 (k_cat_/K_M_ = 277 and 300 *mM*^*-1*^ *s*^*-1*^, respectively). Notably, hCatK displayed between 58 to 63-fold higher catalytic efficiency for FCS-Wu compared to hCatS and hCatL (Fig. 2B,C). Also, hCatK showed 24- and 137-fold higher catalytic efficiency for FCS-Omic compared to hCatL and hCatS. For the TCS substrate, hCatB hydrolyzed the site with ∼1.9-fold higher catalytic efficiency than TMPRSS2, whereas hCatK remained a competent enzyme but operated at ∼3.6-fold lower efficiency than TMPRSS2. Furin and CatL exhibited only trace activities toward TCS (Fig. 2D).

These data indicate that among cysteine cathepsins, hCatB and hCatK display substantial catalytic capacity for both FCS and TCS substrates, in some cases approaching or exceeding the activity of TMPRSS2. Given the exclusive reliance of endosomal viral entry on cathepsins, tissues with high CatB and CatK expression are likely to use these enzymes as dominant endosomal proteases that activate the Spike protein and thereby facilitate viral entry.

### LC-MS/MS analysis of protease cleavage sites using the substrate representing the TCS and FCS from Wuhan and Omicron SARS-CoV-2

Proteolytic activation of the spike protein at FCS and TCS is a critical step in deriving membrane fusion events required for SARS-CoV-2 infectivity^31^. Although multiple host proteases have been implicated, their exact cleavage preferences at the FCS and TCS remain unclear. Defining these patterns is essential to assign enzymatic activity unambiguously and distinguish overlapping or alternative processing events. We therefore performed LC–MS/MS analysis using FRET-based synthetic peptide substrates corresponding to FCS-Wu, FCS-Omic, and TCS. To confirm the cleavage patterns observed with synthetic peptides, we performed complementary LC-MS/MS analysis using full-length recombinant S1 and S2 subunits of the Wuhan-type spike protein as substrates for selected proteases.

LC-MS/MS analysis of the TCS substrate revealed that cleavage after `PS`↓ (Ac-`AE` (E) `LPDPSKPS`↓`KR`) is a unique and preferred site for hCatK (Fig. 3A, B). This cleavage was also confirmed using the S2 subunit as a substrate for hCatK (`QILPDPSKPS`; Fig. 3C, D). Human CatB generated two additional cleavage products after `RS`↓ and `SF`↓ (Ac-`AE` (E) `LPDPSKPSKRS`↓`F`↓`I`) in addition to the canonical hCatB cleavage at `KR`↓ (Fig. 3A). In the S2 subunit analysis the `RS`↓ cleavage of hCatB was also confirmed. TMPRSS2 produced two expected cleavage sites: a major canonical `KR`↓ and a noncanonical `SK`↓ site (Ac-`AE` (E) `LPDPSKPSK`↓`R`↓`SFI`). In contrast, hCatL, consistent with its poor proteolytic efficiency, generated only a few detectable fragments after 2 hours of reaction.

**Figure 3:**
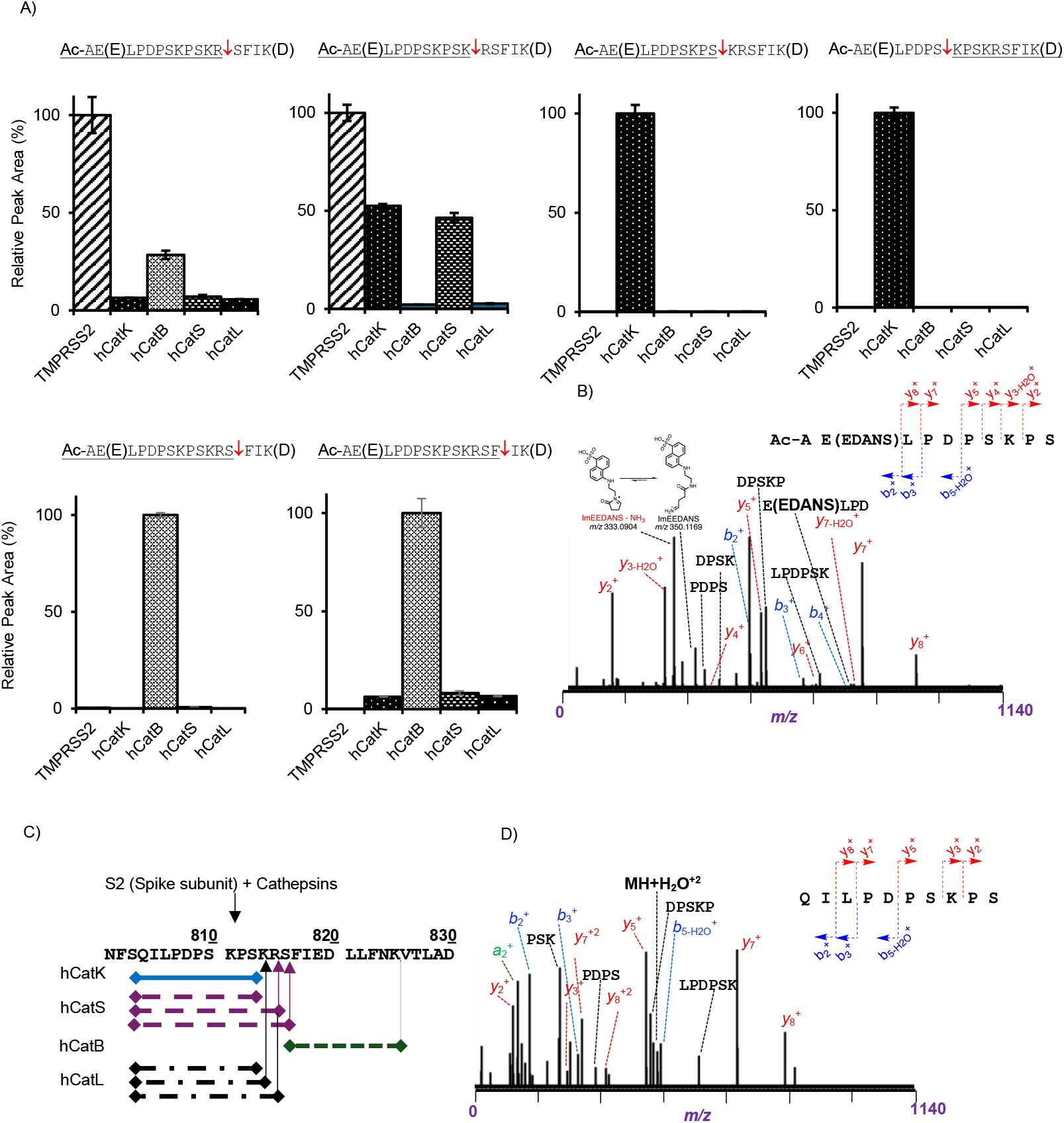
Cleavage site analysis in TMPRSS2 sensitive areas of Spike protein derived substrates. (A) Comparing the relative peak area (%) of proteases cleavage products using TCS as a substrate. (B) The tandem mass spectrum of a selected product peptide (Ac-`AE`(Edans)`LPPSKPS`. (C) Comparing proteases cleavage products in S2’ region using recombinant spike protein as a substrate. (D) The tandem mass spectrum of a selected product peptide `QILPDPSKPS`. Peptides are shown in their full-length sequences; underlined regions indicate fragments that were detected and confirmed by LC-MS/MS. The data were obtained from two independent replicates. Values are reported as mean ± standard deviation (SD).

A similar analysis was performed using the FCS-Wu and FCS-Omic substrates. Notably, hCatK exhibited a newly acquired preference for cleavage at mutation-associated sites within the FCS-Omic sequence compared to FCS-Wu, particularly at Ac-`AE`(E)`QTKSH`↓`RRARSVK`(D) and Ac-`AE`(E)`QTKSHR`↓`RARSVK`(D) (Fig. 4A,B). Furin generated two major cleavage events in both substrates: the canonical `RRAR`↓ site and an additional, less expected cleavage at `Q`↓`TNS` (Ac-`AE`(E)`Q`↓`TNSPRRAR`↓`SVK`(D). Importantly, Furin, consistent with its kinetic parameters, did not exhibit any additional cleavage preferences in Omicron compared to Wuhan (Fig. 4A, C). hCatB cleaved both FCS-Wu and FCS-Omic with a similar efficacy, without a change in substrate cleavage site preference (Fig. 4A, B; Fig. S1). TMPRSS2 produced cleavage after `RR`↓, which overlapped with Furin (at ∼50% lower intensity) and hCatB (at ∼80% lower intensity). In contrast, hCatL showed minimal activity toward both FCS substrates, consistent with its poor kinetic parameters. In the S2-specific analysis, no detectable peptide fragments corresponding to the FCS were observed in the hCatL-catalyzed reactions. These results underscore hCatK and hCatB as proteases with substantial capacity to cleave both TCS and FCS sites, in contrast to the restricted specificity of TMPRSS2 and the minimal activity of hCatL. They also show that Furin and TMPRSS2 did not acquire new cleavage site preferences when comparing Wuhan and Omicron sequences. This suggests that the altered cell entry behavior of Omicron relative to Wuhan is more likely driven by a shift toward endosomal entry pathways, with a greater reliance on cathepsin-mediated activation.

**Figure 4:**
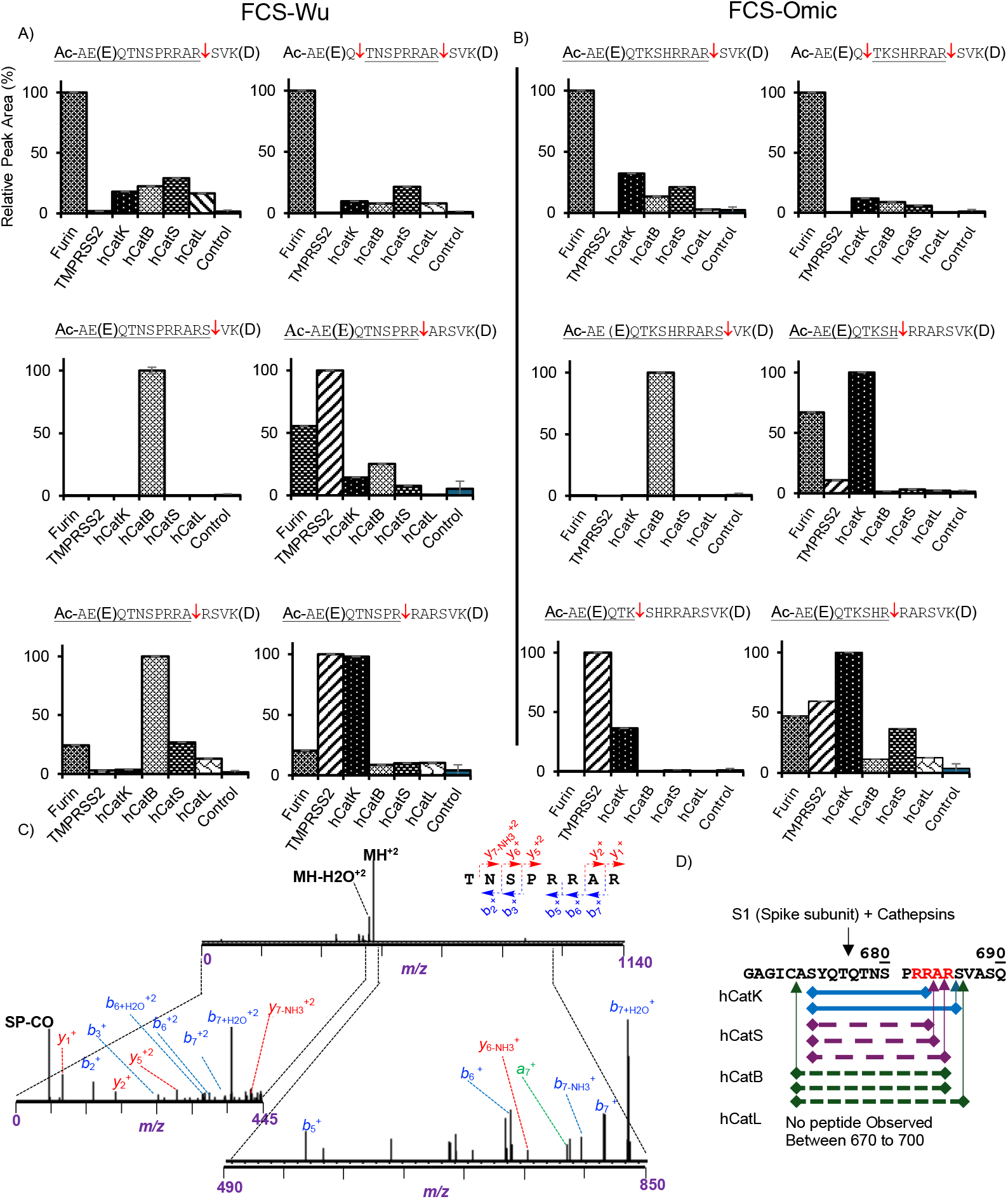
Cleavage site analysis in Furin sensitive areas of Spike protein derived substrates. Comparing the relative peak area (%) of proteases cleavage products using FCS-Wu (A) and FCS-Omic (B) as a substrate. (C) The tandem mass spectrum of a selected product peptide `TNSPRRAR`. (D) Comparing proteases cleavage products in S1/S2 region using recombinant spike protein as a substrate. Peptides are shown in their full-length sequences; underlined regions indicate fragments that were detected and confirmed by LC-MS/MS. The data were obtained from two independent replicates. Values are reported as mean ± standard deviation (SD).

### Cathepsin substrate specificity towards 3CL-Mpro substrate

3CL-Mpro exhibits specificity for cleavage at the (`LQ`↓) motifs, which was the basis for the design of Nirmatrelvir.^17-19^ Notably, `LQ` is also a well-recognized P2 and P1 cleavage site for cathepsins based on the MEROPS database. To assess the substrate specificity of cathepsins in comparison with 3CL-Mpro, we evaluated the enzymatic kinetics of cathepsins using a 3CL-Mpro fluorogenic substrate MCA-`AVLQ`↓`SGFR`(`K`(Dnp)`RR`, and compared the kinetic parameters with those obtained for 3CL-Mpro. The results showed that hCatK (*k*_cat_/K_M_ = 135 *mM*^*-1*^ *s*^*-1*^) and hCatS (*k*_cat_/K_M_ = 133 *mM*^*-1*^ *s*^*-1*^) are about 8-fold more efficient than the viral protease (*k*_cat_/K_M_ = 17.1 *mM*^*-1*^ *s*^*-1*^). In contrast, hCatL and hCatB did not show any significant hydrolytic activity against the 3CL-Mpro substrate. LC-MS/MS analysis of protease cleavage sites showed that, unlike hCatL which showed trace proteolytic activity, hCatK and hCatS efficiently cleaved the 3CL-Mpro substrate at `LQ`↓. In addition, hCatK also showed an expected cleavage site after `FR`↓ (MCA-`AVLQ`↓`SGFR`↓(`K`(Dnp)`RR`) (Fig. 5B,C,D). These findings demonstrate that both hCatK and hCatS have a strong capacity to recognize and hydrolyze the canonical 3CL-Mpro substrate at the `LQ`↓ motif with an efficiency substantially exceeding that of the viral protease itself.

**Figure 5:**
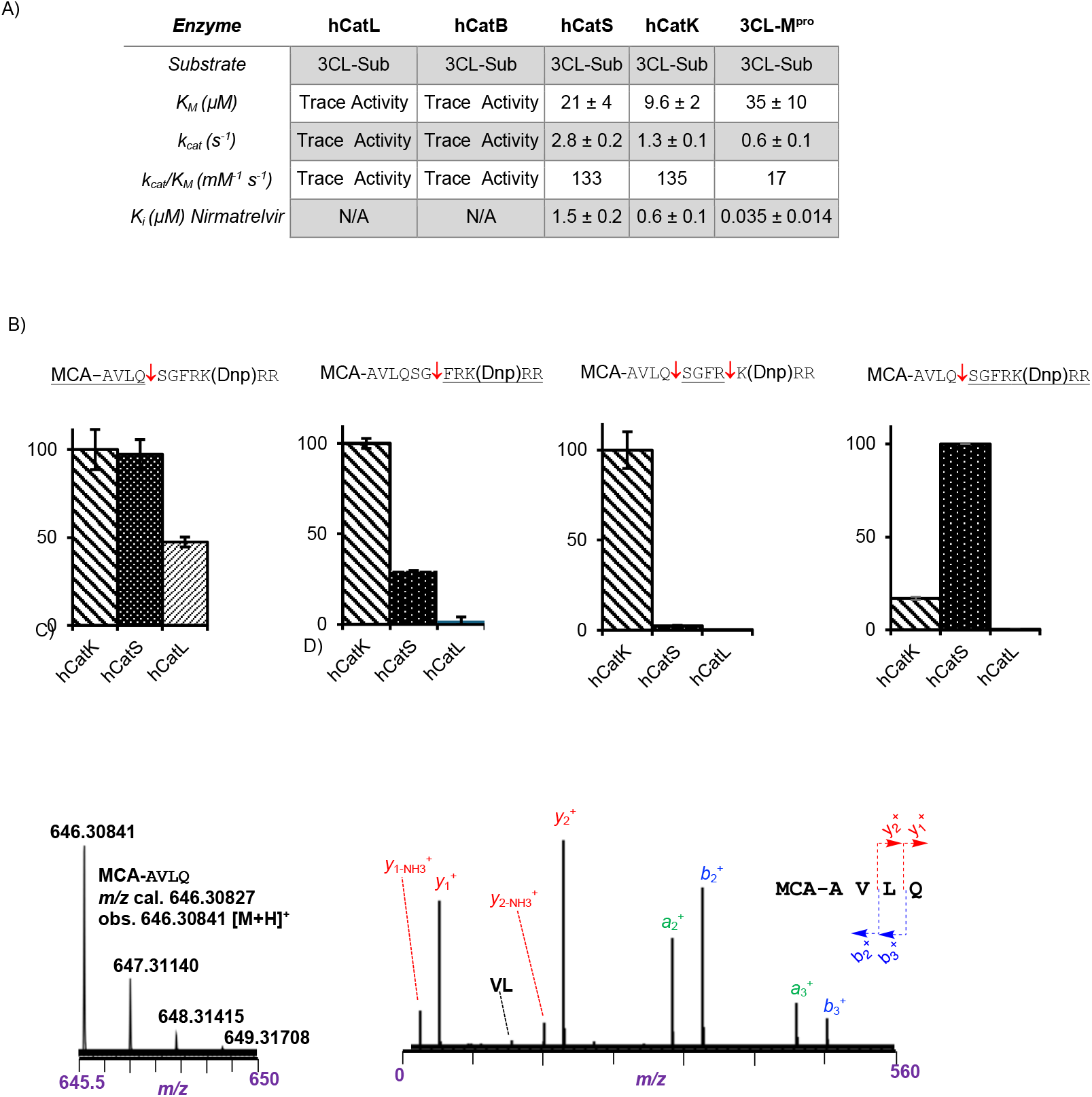
Kinetic and cleavage site analysis of human cathepsins and 3CL-Mpro of a substrate representing the NSP4-NSP5 processing site and their K_i_ values towards Nirmatrelvir. (A) Kinetic parameters of cathepsins and 3CL-Mpro using a standard 3CL-Mpro substrate (MCA-`AVLQSGFRK`(Dnp)`RR`). (B) Comparing the relative peak area (%) of cathepsins cleavage products using 3CL-Mpro substrate. (C) mass spectrum of MCA-`AVLQ` which a major product of 3CL-Mpro and hCatK and hCatS. (D) tandem mass spectrum of MCA-`AVLQ`. Peptides are shown in their full-length sequences; underlined regions indicate fragments that were detected and confirmed by LC-MS/MS. The kinetic data were obtained from two independent experiments, each performed in triplicate. Values are reported as mean ± standard deviation (SD).

### Interaction of Nirmatrelvir with cathepsins

Given that hCatK, hCatS, and 3CL-Mpro are efficient to cleave substrates at the `LQ`↓ motif, we hypothesized that they might also share susceptibility to Nirmatrelvir, the inhibitor specifically designed for 3CL-Mpro^19^. To evaluate the dual inhibitory potential of Nirmatrelvir, we determined the K_i_ values for cathepsins. Nirmatrelvir inhibited hCatK (K_i_ = 600 nM) with the highest efficacy among the cathepsins tested. Cathepsin S showed an intermediate inhibition with 1.5 µM. hCatB and hCatL were not inhibited by 10 µM Nirmatrelvir (Fig. 5A). These results, together with the kinetic data and the Odanacatib-induced viral proliferation inhibition, indicate that hCatK and 3CL-Mpro represent promising targets for dual inhibition by Nirmatrelvir.

### Crystal structure of hCatK/Nirmatrelvir complexes

The submicromolar inhibitory constant of Nirmatrelvir against hCatK may serve as a basis for designing more effective dual inhibitors targeting both 3CL-Mpro and hCatK. To understand how Nirmatrelvir inhibits hCatK, we determined the co-crystal structure of hCatK following covalent adduct formation with Nirmatrelvir at 1.9 Å resolution (Table S1). Nirmatrelvir binds to the active site of hCatK in a manner similar to Odanacatib, forming a covalent bond with the sulfur atom of the active site Cys25 and the nitrile group of the inhibitors (Fig. 6.C & D, Fig. S2). It also forms comparable hydrogen bonds with residues Gly66 and Asn161. The hydrogen bond between Nirmatrelvir and the oxyanion hole residue, Gln19 of hCatK is unique when compared to the hCatK/Odanacatib complex. Well-ordered water molecules were observed in proximity to the enzyme’s active site, interacting with the inhibitor. These water molecules form a hydrogen bond network involving the nitrile group, oxygen of pyrrolidine ring, and the oxygen near the 3-azabicyclo[3.1.0]hexane ring of Nirmatrelvir (Fig. 6A, shown in red), suggesting a potential role in stabilizing the inhibitor binding (Fig. 6.D). The active site of apo hCatK (PDB ID 5TUN) was aligned with the Nirmatrelvir-bound hCatK structure. Overall, the active sites appear highly similar, except for Tyr67, which is rotated 17.8° upward toward Nirmatrelvir (Fig. 6F). However, unlike Odanacatib, Nirmatrelvir lacks the π–π interaction between Tyr67 and a biphenyl ring, a key interaction present in the hCatK:Odanacatib adduct co-crystal structure (PDB 5TDI; Fig. 6G). This absence may explain the higher potency of Odanacatib, which exhibits a subnanomolar inhibition constant for hCatK (K_i_ = 0.18 ± 0.06 nM)^14^. Structural comparison of hCatK in its apo form and in complex with either Odanacatib or Nirmatrelvir reveals no significant conformational changes, with an RMSD of ∼0.2 Å (Fig. S1; superposition was done with the all atoms), indicating that inhibitor binding does not induce major structural rearrangements. These findings suggest that incorporating a phenyl ring into the Nirmatrelvir scaffold to enable π–π stacking with Tyr67 could enhance its binding affinity and inhibitory potency toward hCatK without interfering with its binding to 3CL-Mpro.

**Figure 6.**
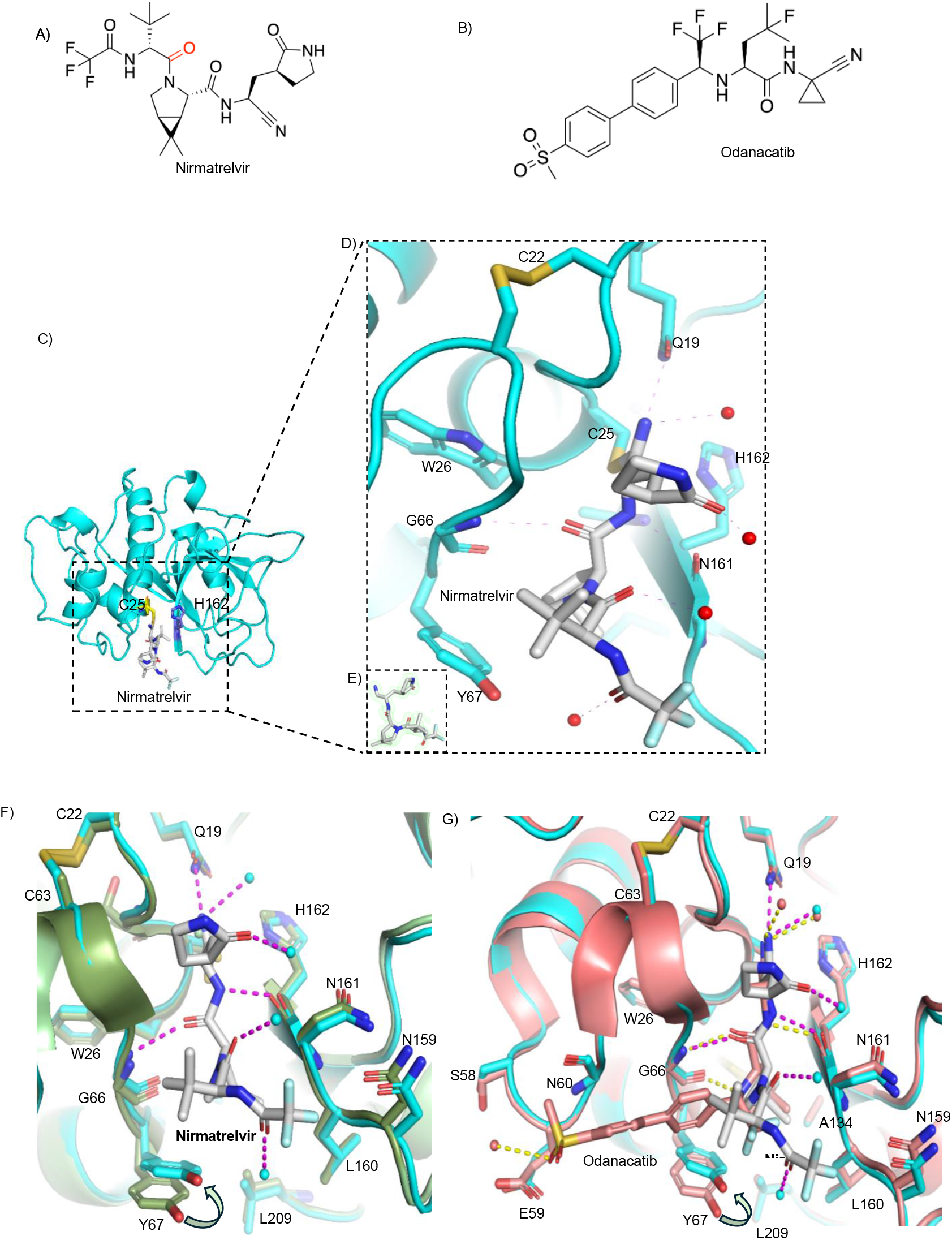
Crystal structure of hCatK with Nirmatrelvir. Structures of A) Nirmatrelvir and (B) Odanacatib (C) Nirmatrelvir bound at the active site of hCatK. (D) Polar interaction of Nirmatrelvir with active site amino acids (*cyan*) and water (*red*) shown in *magenta*. (E) Omit map of Nirmatrelvir contoured at σ 3. (F) Comparison of the active site of hCatK in the presence and absence of Nirmatrelvir. Uninhibited hCatK shown in *palegreen*; Nirmatrelvir (white colour) bound hCatK and water molecules were shown in *cyan*. (G) Comparison of active site of hCatK bound to Odanacatib and Nirmtrelvir. Odanacatib bound hCatK and water molecules were shown in *deepsalmon;* Polar interaction of Odanacatib are shown in *yellow*.

## Discussion

This study defines hCatK as a previously underappreciated yet catalytically dominant host protease that drives SARS-CoV-2 spike activation and reveals its high-affinity interaction with the FDA-approved 3CL-Mpro inhibitor nirmatrelvir. In contrast to earlier reports implicating hCatL as the main endosomal facilitator of spike processing^6^, our data demonstrate that hCatK exhibits markedly greater catalytic and functional activity. The supporting evidence for hCatL in prior studies largely derived from inhibition assays using small molecule inhibitors, yet most inhibitors used were broad-spectrum cysteine protease blockers of limited protease specificity such as E-64d and MDL-28170^6,32^. Moreover, the more selective hCatL inhibitor SID26681509 displayed inhibitory efficacy nearly tenfold weaker than the pan-cysteine inhibitor E-64d at concentration of 10 µM^6^. By contrast, the hCatK-selective inhibitor Odanacatib suppressed viral propagation with a potency equivalent to E-64d at 10 µM, providing direct functional evidence for hCatK as a critical catalytic driver of spike activation.

Enzyme activity profiling revealed that hCatK cleaves spike FCS with 24-to 63-fold higher efficiencies compared to hCatL. Although Cathepsin B exhibits detectable cleavage of spike peptides in vitro, inhibition of this enzyme does not measurably affect viral entry, suggesting that its proteolytic activity is not functionally utilized during infection^7^.

While catalytic efficiency of hCatK were similar for Wuhan and Omicron FCS substrates, LC–MS/MS mapping revealed distinct cleavage signatures. A unique Omicron-specific cleavage site preferentially processed by hCatK suggests that sequence adaptations within variant spikes may modulate protease utilization and fusion dynamics. This observation aligns with reports of enhanced reliance on cathepsin-dependent entry in Omicron, reinforcing the importance of defining exact proteolytic events in the S2′ region adjacent to the fusion peptide, determinants that govern membrane fusion efficiency^4^. For instance, the T813S substitution (Ac-`AE`(E)LPDPSKP**<u>S</u>**KRSFIK(D)) that arose during SARS-to-SARS-CoV-2 evolution introduces an additional S2′ cleavage motif that we find preferentially processed by hCatK, implicating the key role of this enzyme in spike fusogenicity and syncytia formation^33^.

Remarkably, hCatK exhibited approximately eight-fold higher catalytic efficiency than 3CL-Mpro when tested on a FRET substrate spanning the NSP4-NSP5 junction (`LQ`↓`SG`). LC–MS/MS analysis confirmed cleavage at the canonical `LQ` site, challenging previous assumptions that this motif was unique to 3CL-Mpro^19^. Nirmatrelvir inhibited hCatK at submicromolar concentrations (K_i_ = 600 nM). This finding is consistent with an earlier reported K_i_ value of 231 nM^34^. The capacity of hCatK to accommodate the `LQ`↓`X` motif within its active site pocket, rationalizes the observed cross-reactivity with Nirmatrelvir, a covalent nitrile inhibitor originally optimized for 3CL-Mpro. Our co-crystal structure of the hCatK–Nirmatrelvir complex supports this mechanism and defines the molecular determinants that enable dual inhibition across viral and host proteases.

Given the restricted expression of hCatK but its enrichment in infection-relevant lung epithelial and vascular endothelial cells targeting hCatK may provide therapeutic selectivity with limited systemic toxicity^8,9^. A dual-inhibition strategy that concurrently engages hCatK and 3CL-Mpro could therefore block both spike activation and viral replication, mitigating resistance risks inherent to single-agent therapies. Such a framework is particularly pertinent for variants like Omicron, which exhibit increased dependence on cathepsin-mediated activation. Because hCatK inhibition in antiviral contexts would be transient, the adverse skeletal effects associated with chronic anti-resorptive therapy are unlikely to pose clinical concern during acute infection^35,36^.

While our findings provide integrated biochemical, kinetic, and mechanistic evidence implicating hCatK in SARS-CoV-2 spike activation, further validation in physiologically relevant *in vivo* models is warranted. Future work should delineate tissue-specific protease expression and viral tropism to refine the therapeutic window for hCatK-directed intervention. Moreover, because other viruses such as Ebola also depend on cathepsin-mediated glycoprotein priming, the structural framework described here may guide rational development of broad-spectrum or dual-acting protease inhibitors^37^. Collectively, our results reposition hCatK, not hCatL, as a central host protease in SARS-CoV-2 pathogenesis and establish a chemical and structural foundation for designing next-generation dual inhibitors targeting both viral and host cysteine proteases.

## Author contributions

SYTD, DS, PP, and DB designed the study, validated results, and were responsible for the administration of project. Specifically, SYTD performed substrate kinetics and all LC-MS/MS analyses and OH executed sample preparation for LC-MS/MS. DS also performed substrate kinetics and was responsible together with FvP and YSC for the crystallization and structural analysis of the CatK/Nirmatrelvir complex. MN performed all BSL3 SARS-CoV-2 endothelial infection experiments and PP was responsible for all cell-based analyses. EM produced and purified recombinant human CatK. SYTD produced and purified human CatS. DS and SYTD produced and purified SARS-CoV-2 3CL-Mpro. BJF and CA produced recombinant TMPRSS2. SYTD, DS, and DB wrote the original draft of the manuscript. All authors reviewed and revised the final manuscript. DB acquired funding for the study.

## Acknowledgements

This work was supported by Canadian Institutes of Health Research grants (PJT-155979, CPG-158275), Collaborative Health Research Project (NSERC CHRP 523434-18) and the NSERC discovery grant (LJGP GR003266). DB was supported by Canada Research Chair award funding. We are thankful to the proteomics core facility at UBC under the leadership of L. Foster and the technical support by J. Rogalski to generate the raw LC-MS/MS data.

## References

1. Shang, J. et al. Cell entry mechanisms of SARS-CoV-2. Proceedings of the National Academy of Sciences of the United States of America 117, 11727–11734 (2020).

2. Shapira T. M.I., Dion SP, Buchholz DW, Imbiakha B, Olmstead AD, Jager M, Désilets A, Gao G, Martins M, Vandal T. A TMPRSS2 inhibitor acts as a pan-SARS-CoV-2 prophylactic and therapeutic. nature 605, 340–348 (2022).

3. Papa G. M.D., Albecka A, Welch LG, Cattin-Ortolá J, Luptak J, Paul D, McMahon HT, Goodfellow IG, Carter A, Munro S. Furin cleavage of SARS-CoV-2 Spike promotes but is not essential for infection and cell-cell fusion. Plos Pathogens 17, e1009246 (2021).

4. Willett, B.J. et al. SARS-CoV-2 Omicron is an immune escape variant with an altered cell entry pathway (vol 7, pg 1161, 2022). Nature Microbiology 7, 1709–1709 (2022).

5. Meng, B. et al. Altered TMPRSS2 usage by SARS-CoV-2 Omicron impacts infectivity and fusogenicity. Nature 603, 706– (2022).

6. Zhao, M.M. et al. Cathepsin L plays a key role in SARS-CoV-2 infection in humans and humanized mice and is a promising target for new drug development. Signal Transduction and Targeted Therapy 6(2021).

7. Ou, X.Y. et al. Characterization of spike glycoprotein of SARS-CoV-2 on virus entry and its immune cross-reactivity with SARS-CoV. Nature Communications 11(2020).

8. Bühling, F. et al. Expression of cathepsin K in lung epithelial cells. Am J Respir Crit Care Med 20, 612–619 (1999).

9. Platt, M.O. et al. Expression of cathepsin K is regulated by shear stress in cultured endothelial cells and is increased in endothelium in human atherosclerosis. American Journal of Physiology-Heart and Circulatory Physiology 292, H1479–H1486 (2007).

10. Bühling, F. et al. Expression of cathepsins B, H, K, L, and S during human fetal lung development. Developmental Dynamics 225, 14–21 (2002).

11. Riva, L. et al. Discovery of SARS-CoV-2 antiviral drugs through large-scale compound repurposing. Nature 586, 113–119 (2020).

12. Ma, X.Y.R. et al. MPI8 is Potent against SARS-CoV-2 by Inhibiting Dually and Selectively the SARS-CoV-2 Main Protease and the Host Cathepsin L**. Chemmedchem 17(2022).

13. Costanzi, E. et al. Structural and Biochemical Analysis of the Dual Inhibition of MG-132 against SARS-CoV-2 Main Protease (Mpro/3CLpro) and Human Cathepsin-L. International Journal of Molecular Sciences 22(2021).

14. Law, S. et al. Identification of mouse cathepsin K structural elements that regulate the potency of odanacatib. Biochemical Journal 474, 851–864 (2017).

15. Gauthier, J.Y. et al. The discovery of odanacatib (MK-0822), a selective inhibitor of cathepsin K. Bioorganic & Medicinal Chemistry Letters 18, 923–928 (2008).

16. Rawlings, N.D. et al. The database of proteolytic enzymes, their substrates and inhibitors in 2017 and a comparison with peptidases in the PANTHER database. Nucleic Acids Research 46, D624–D632 (2018).

17. Yaghi, R.M., Andrews, C.L., Wylie, D.C. & Iverson, B.L. High-Resolution Substrate Specificity Profiling of SARS-CoV-2 Mpro; Comparison to SARS-CoV Mpro. Acs Chemical Biology 19, 1474–1483 (2024).

18. Chuck, C.P. et al. Profiling of Substrate Specificity of SARS-CoV 3CL. Plos One 5(2010).

19. Owen, D.R. et al. An oral SARS-CoV-2 M(pro) inhibitor clinical candidate for the treatment of COVID-19. Science 374, 1586–+ (2021).

20. Linnevers, C.J. et al. Expression of human cathepsin K in Pichia pastoris and preliminary crystallographic studies of an inhibitor complex. Protein Science 6, 919–921 (1997).

21. Andrault, P.M., Panwar, P. & Bromme, D. Characterization of cathepsin S exosites that govern its elastolytic activity. Biochem J 477, 227–242 (2020).

22. Fraser, B.J. et al. Structure and activity of human TMPRSS2 protease implicated in SARS-CoV-2 activation. Nature Chemical Biology 18, 963–+ (2022).

23. Battye, T.G., Kontogiannis, L., Johnson, O., Powell, H.R. & Leslie, A.G. iMOSFLM: a new graphical interface for diffraction-image processing with MOSFLM. Acta Crystallogr D Biol Crystallogr 67, 271–81 (2011).

24. Winn, M.D. et al. Overview of the CCP4 suite and current developments. Acta Crystallogr D Biol Crystallogr 67, 235–42 (2011).

25. Moriarty, N.W., Grosse-Kunstleve, R.W. & Adams, P.D. Electronic Ligand Builder and Optimization Workbench (eLBOW): a tool for ligand coordinate and restraint generation. Acta Crystallographica Section D-Structural Biology 65, 1074–1080 (2009).

26. Adams, P.D. et al. PHENIX: a comprehensive Python-based system for macromolecular structure solution. Acta Crystallogr D Biol Crystallogr 66, 213–21 (2010).

27. Emsley, P., Lohkamp, B., Scott, W.G. & Cowtan, K. Features and development of Coot. Acta Crystallographica Section D-Biological Crystallography 66, 486–501 (2010).

28. Matarese, A., Gambardella, J., Sardu, C. & Santulli, G. miR-98 Regulates TMPRSS2 Expression in Human Endothelial Cells: Key Implications for COVID-19. Biomedicines 8(2020).

29. Wagner, J.U.G. et al. Increased susceptibility of human endothelial cells to infections by SARS-CoV-2 variants. Basic Research in Cardiology 116(2021).

30. Hoffmann, M. et al. Camostat mesylate inhibits SARS-CoV-2 activation by TMPRSS2-related proteases and its metabolite GBPA exerts antiviral activity. Ebiomedicine 65(2021).

31. Lavie, M., Dubuisson, J. & Belouzard, S. SARS-CoV-2 Spike Furin Cleavage Site and S2′ Basic Residues Modulate the Entry Process in a Host Cell-Dependent Manner. Journal of Virology 96(2022).

32. Simmons, G. et al. Inhibitors of cathepsin L prevent severe acute respiratory syndrome coronavirus entry. Proceedings of the National Academy of Sciences of the United States of America 102, 11876–11881 (2005).

33. Ma, Y. et al. Spike substitution T813S increases Sarbecovirus fusogenicity by enhancing the usage of TMPRSS2. Plos Pathogens 19(2023).

34. Duveau, D.Y. & Thomas, C.J. The Remarkable Selectivity of Nirmatrelvir. Acs Pharmacology & Translational Science 5, 445–447 (2022).

35. Rünger, T.M. et al. Morphea-like skin reactions in patients treated with the cathepsin K inhibitor balicatib. Journal of the American Academy of Dermatology 66, E89–E96 (2012).

36. Mullard, A. Merck & Co. drops osteoporosis drug odanacatib. Nature Reviews Drug Discovery 15, 669–669 (2016).

37. Bestle, D. et al. Novel proteolytic activation of Ebolavirus glycoprotein GP by TMPRSS2 and cathepsin L at an uncharted position can compensate for furin cleavage. Virus Research 347(2024).

