## Supplementary material for "Cathepsin K as a Key Protease in Processing of SARS-CoV-2 Spike Activation Sites and a Target of Dual-Inhibition": Supplimentary info

[1]Department of Oral Biological & Medical Sciences, Faculty of Dentistry, The University of British Columbia; Vancouver, Canada. [2] Department of Biochemistry & Molecular Biology, Faculty of Medicine, The University of British Columbia; Vancouver, Canada. [3]Faculty of Health Sciences, Simon Fraser University, Burnaby, Canada. [4] Department of Medical Biophysics, University of Toronto, Toronto, Canada. [5]Structural Genomics Consortium Toronto, Toronto, Canada.

[#]these authors contributed equally

**Table S1.** Statistics of data collection and refinement

| | hCatK + Nirmatrelvir |
|---|---|
| **Data collection** | |
| X-ray source | SSRL BL12-2 |
| Wavelength (Å) | 0.97946 |
| Space group | $P4_32_12$ |
| Cell dimensions | |
| *a*, *b*, *c* (Å) | 61.77, 61.77, 111.93 |
| *α, β, γ* (°) | 90.0, 90.0, 90.0 |
| Resolution (Å) | 54.08–1.64 (1.67–1.64) |
| Redundancy | 7.7 (3.5) |
| Completeness (%) | 84.3 (41.7) |
| $I/\sigma I$ | 13.3 (1.3) |
| $CC_{1/2}$ | 0.998 (0.430) |
| **Refinement** | |
| Resolution (Å) | 43.67 – 1.90 |
| No. of reflections | 17506 |
| $R_{work}/R_{free}$ | 0.163/0.212 |
| No. of residues | |
| Protein | 215 |
| Ligands | 3 |
| Water | 245 |
| *B*-factors | |
| Protein | 14.2 |
| Ligands | 22.1 |
| Water | 24.7 |
| RMSDs | |
| Bond lengths (Å) | 0.002 |
| Bond angles (°) | 0.52 |
| Ramachandran plots | |
| Favored (%) | 96.7 |
| Allowed (%) | 3.3 |
| Disallowed (%) | 0.0 |
| **PDB code** | 9O69 |

[1]Highest resolution shell is shown in parentheses

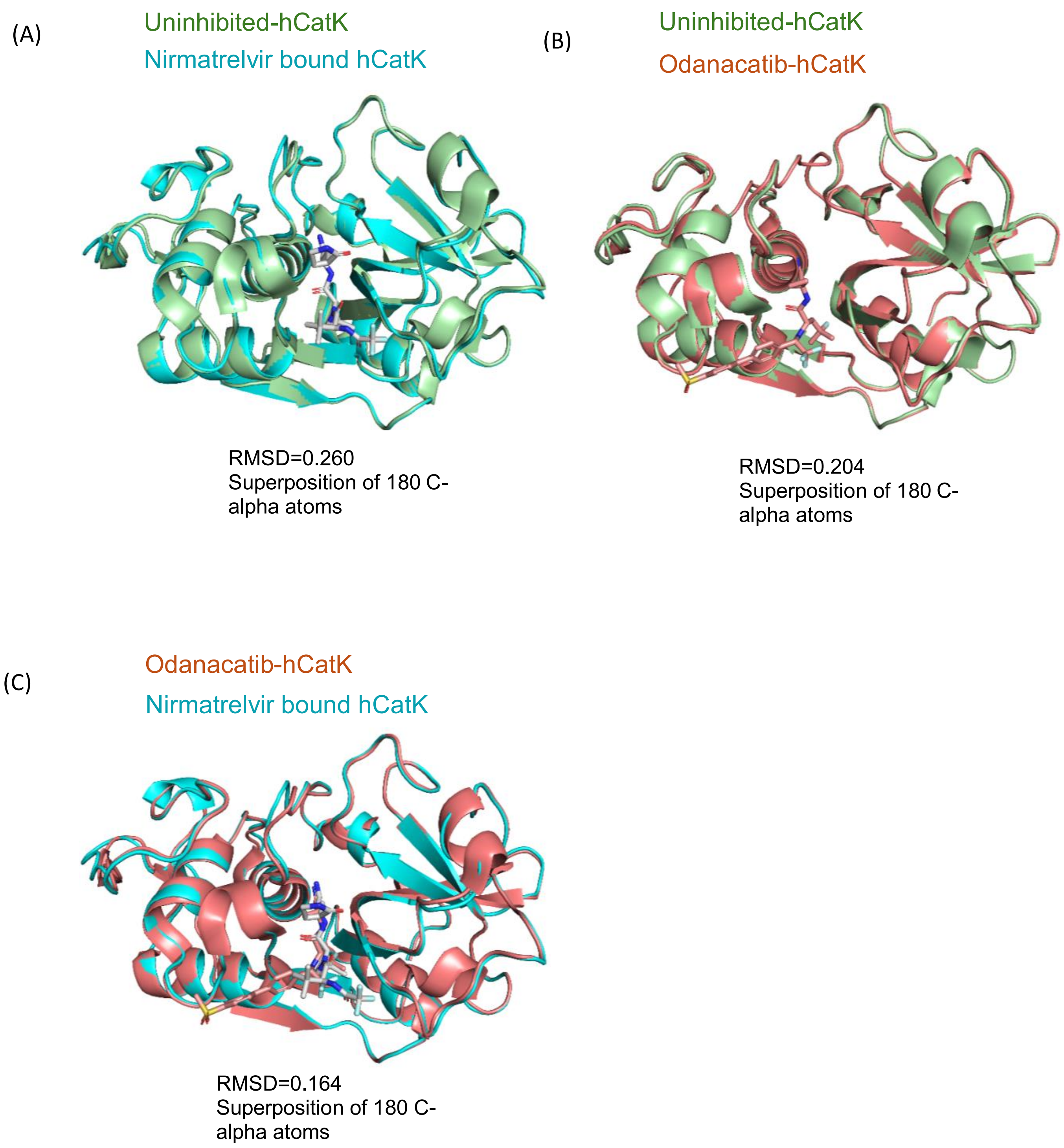


**Fig. S1**. **Secondary structure comparisons between free hCatK and CatK inhibitor complexes**. **(**A) alignment of uninhibited hCatK (5TUN; *palegreen*) with Nirmatrelvir bound hCatK (this study; *cyan*). (B) alignment of uninhibited hCatK (5TUN) with ODN bound hCatK (5TDI; *deepsalmon*). (C) alignment of ODN bound hCatK (5TDI) with Nirmatrelvir bound hCatK (this study; *cyan*).

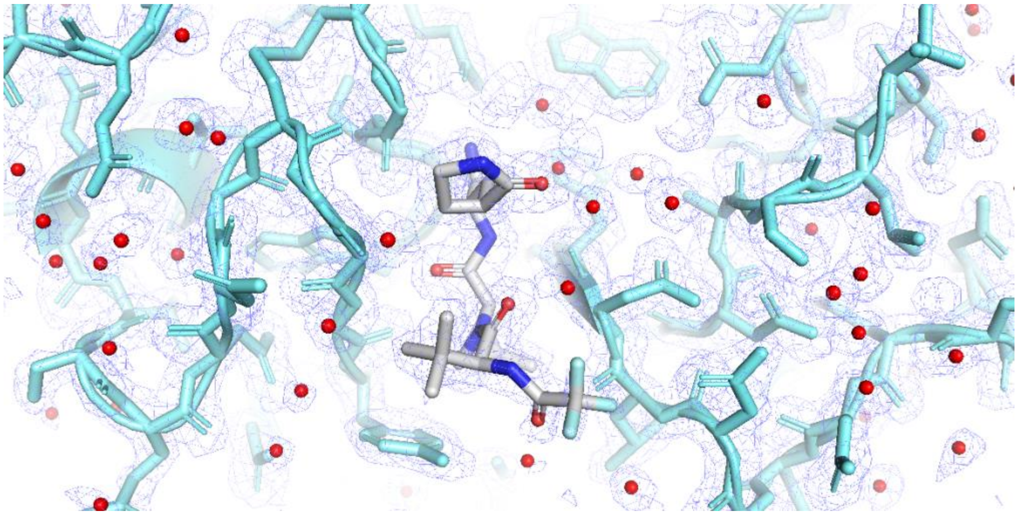

**Fig. S2**. Structures of Nirmatrelvir (white) bound at the active site of the hCatK (cyan). Electron density map contoured at σ 1.
